# Human posterior insula to posterior cingulate functional connectivity is related to the subjective intensity of acute pain

**DOI:** 10.1101/246959

**Authors:** Christopher J. Becker, Keith M. Vogt, James W. Ibinson

## Abstract

**Background:** Several studies have shown that, in the context of chronic pain, alterations in functional connectivity between the insula and default mode network (DMN) are correlated with pain intensity. However, it is unknown whether a similar relationship between insula-to-DMN connectivity and pain intensity exists in the absence of chronic pain. Our aims were 1) to assess whether insula-to-DMN connectivity changes are correlated to the intensity of ongoing acute pain in healthy subjects, and 2) provide an initial assessment of the capacity of pIns-to-PCC connectivity to differentiate between painful and non-painful states. 3T BOLD functional imaging data were obtained from 13 healthy human subjects in three conditions: rest, non-painful tactile stimulation, and capsaicin-induced pain.

**Results:** Posterior insula to posterior cingulate (pIns-to-PCC) connectivity was significantly correlated with pain intensity. Additionally, we found that discrimination between painful and non-painful states could be achieved based on pIns-to-PCC connectivity.

**Conclusions:** Functional connectivity alterations in healthy subjects experiencing acute ongoing pain are similar to those observed in chronic pain patients, and can be used to classify pain state. This suggests that the inclusion of the pIns and PCC regions in future fcMRI-based methods to detect ongoing pain is warranted.

## Background

Functional connectivity MRI is a technique that relies on correlations in spontaneous, low frequency (<0.1 Hz) fluctuations of the blood-oxygen level dependent (BOLD) signal. These correlations reveal coherent networks of anatomically distinct, yet functionally connected brain regions, even in the absence of experimental stimuli (Biswal et al. 1995; Fox et al. 2005). One prominent example is the default mode network (DMN), which typically consists of brain structures that show greater connectivity at rest than during task performance, and is thought to be involved in self-referential thinking and retrieval of autobiographical memory (Buckner et al. 2008; Mak et al. 2016). Both acute and chronic pain are associated with alterations in connectivity involving the DMN (Baliki et al. 2014; Mantini et al. 2009), particularly the posterior cingulate cortex (PCC), which is a central hub within the DMN (Fransson and Marrelec 2008; Leech et al. 2012).

The insula is an important area for pain processing (Apkarian et al. 2005; Garcia-Larrea 2012; Segerdahl et al. 2015). Connectivity between the insula and DMN has been shown to be characteristically altered by chronic pain (Baliki et al. 2014). Specifically, in chronic pain patients, insula-to-DMN connectivity is increased in proportion to pain intensity (Baliki et al. 2014; Napadow et al. 2010), with pain-exacerbating maneuvers further increasing insula-to-DMN connectivity (Loggia et al. 2013), and pain relief leading to decreased connectivity (Ceko et al. 2015; Napadow et al. 2012). Taken together, these findings suggest that, in the context of chronic pain, alterations in insula-to-DMN connectivity track changes in pain intensity.

In the absence of chronic pain, however, it is unclear whether such a relationship exists. We previously demonstrated that posterior insula (pIns) to PCC connectivity is selectively altered (as opposed to the anterior insula) in response to acute pain created by electrical nerve stimulation in healthy subjects (Vogt et al. 2016). It is unknown, however, whether the magnitude of connectivity change is related to the intensity in acute pain. Furthermore, it is unclear whether previously observed pIns-to-PCC connectivity changes are pain-modality specific, or would be seen in response to any salient stimulus.

The primary aim of this study was to investigate whether posterior insula-to-DMN connectivity is altered in proportion to pain intensity in healthy subjects when using capsaicin as source of acute pain. To do this, we compared whole-brain functional connectivity during ongoing acute pain to connectivity at rest, as well as during a salient non-painful control condition. We focused specifically on pIns-to-PCC connectivity. Our hypotheses were 1) pIns-to-PCC connectivity would be altered in proportion to pain intensity, and 2) this connectivity would not be altered by salient, but non-painful sensory stimulation. A secondary aim was to provide an initial assessment of the capacity of pIns-to-PCC connectivity to discriminate between painful and non-painful states.

## Methods

### Subjects

Institutional Review Board approval was obtained from the University of Pittsburgh, and informed consent was obtained from each subject. A total of 13 right-handed healthy adults (6 male) between the ages of 24-35 participated. Exclusion criteria included diagnosis of a neurologic disease, use of prescription medications or illicit drugs, pregnancy, and any contraindication to MRI. Due to the pilot nature of this study, subjects were enrolled in two cohorts. The first six subjects were in Cohort 1, the next seven in Cohort 2. All data were combined across cohorts for analysis.

### Experimental Conditions

Two scans were obtained from each subject in Cohort 1. The first was a rest scan, in which subjects were instructed to not think of anything in particular and to keep their eyes open. They were not instructed to maintain focus on any particular object (i.e., no fixation cross was projected into the scanner). The second scan occurred after the application of a capsaicin-soaked 1×1 cm gauze pad to the left volar forearm. The capsaicin solution was diluted to a concentration of 0.004 Molar in 70% ethanol, and the pad was covered with waterproof adhesive film. Scans were obtained an average of 25 minutes after the application of capsaicin (range 22-28 minutes).

In Cohort 2, four scans were obtained from each subject. The first was a rest scan, identical to Cohort 1. The second was a touch scan, which was obtained as a researcher continuously moved a gauze pad around the left volar forearm of each subject. Velocity, direction, and pressure were continuously varied to maximize the salience of this stimulus. Subjects were instructed to attend to this tactile stimulus, and were told that they would be asked questions about it after the procedure. After application of capsaicin, which was done in an identical manner to Cohort 1, the third and fourth scans occurred. An initial scan was obtained immediately after the application of capsaicin, and a final scan was obtained once pain ratings had reached at least 5/10. This occurred an average of 18 minutes after application of capsaicin (range 14-23 minutes).

The sensation elicited by the topical application of capsaicin typically begins with a period of “tingling”, followed by increasingly intense pain, the timing of which varies from subject to subject. Thus, the initial set of capsaicin scans from Cohort 2 were expected to capture mostly the tingling aspect, although some would likely experience mild pain, whereas the second set was expected to capture moderate to severe pain in all subjects. Thus, scans were assigned to one of three conditions for analysis: *Rest, Pain*, and *Tactile Sensation*. *Rest* included all of the resting scans without stimulation. *Pain* included all of the capsaicin scans that were associated with a pain rating of 3/10 or greater. *Tactile Sensation* included all of the touch scans, as well as any capsaicin scans that were associated with a pain rating of 2/10 or lower, as this sensation was most commonly described as tingling instead of pain.

### Data Acquisition

Subjective pain ratings were obtained at the end of each capsaicin scan. Subjects responded by verbally rating the intensity of their pain on a scale from 0 to 10, with 0 representing no pain and 10 representing the worst pain imaginable. In the rest and touch conditions, subjects verbally confirmed that they did not experience pain during the scan. Salience ratings for the touch and capsaicin stimuli were also obtained after completion of the procedure by the subjects in Cohort 2. They responded on a scale from 0 to 10, with 0 being “not prominent in your thoughts at all” and 10 being “something that you could not take you mind off”.

The functional scans for Cohort 1 were each six minutes in duration. For Cohort 2, each functional scan lasted eight minutes. For all subjects in both cohorts, a single high-resolution structural image was obtained. Images for Cohort 1 were collected with a 3.0-T Siemens Trio scanner (Siemens Medical Solutions USA, Inc, Malvern, PA). High-resolution structural images were collected for each participant to facilitate the creation of group maps using a T1-weighted scanning technique (MPRAGE sequence, TR/TE/Flip = 1.6 s/3.18 ms/8°; field of view = 256 cm × 240 cm; slice thickness = contiguous 1.2 mm; in-plane resolution = 1.0 mm). Functional images were collected using a BOLD sensitive single-shot gradient echo, T2*- weighted sequence (TR/TE/ Flip = 2 s/30 ms/90°; 64 × 64 matrix; slice thickness = 3 mm, in-plane resolution = 3.125 × 3.125 mm). Twenty-eight axial slices with no gaps were acquired in an interleaved fashion.

Images for Cohort 2 were collected with a Siemens Allegra 3T MRI Head-Scanner. An MPRAGE sequence (TR/TE/Flip = 1.54 s/3.04 ms/8°; field of view = 256 mm × 256 mm; slice thickness = contiguous 1 mm; in-plane resolution = 1 mm; matrix = 256 × 256) was again used for high-resolution scanning. Functional imaging consisted of 38 axial slices collected with BOLD weighted single-shot gradient echo parameters (TR/TE/ Flip = 2 s/25 ms/70°; 64 × 64 matrix; slice thickness = 3.2 mm, in-plane resolution = 3.125 × 3.125 mm).

### Data Analysis

All functional images were preprocessed within MATLAB (Mathworks Inc., Natick, MA, USA) using the Functional Connectivity Toolbox, version 15.e (Whitfield-Gabrieli and Nieto-Castanon 2012). Preprocessing included realignment, slice timing correction, segmentation, normalization, and spatial smoothing using a Gaussian kernel of full width at half maximum of 6 mm. Each participant’s functional images were registered to their high-resolution T1-weighted structural image, and to the MNI152 standard space template. The preprocessed data were band-pass filtered (0.009 < *f* < 0.08 Hz), and motion artifacts were removed by linear regression of six motion parameters obtained by rigid body correction of motion of the head. Grey matter, white matter, and CSF signals were included as regressors of no interest, as this has been shown to improve specificity for connectivity maps of pain (Ibinson et al. 2015).

BOLD signal time courses were extracted from a pre-defined ROI in the right posterior insula, contralateral to the touch and capsaicin stimuli. This ROI was defined as a sphere with a radius of 6 mm centered around MNI coordinates 38, −4, 10, as identified in a previous study of acute electrical nerve stimulation pain (Ibinson and Vogt 2013). The signal was averaged across the voxels within this ROI for each subject, and the resulting time courses were used as explanatory variables in a seed-based analysis. Using procedures established by Fox et al. (Fox et al. 2005), correlation maps were generated by computing the correlation coefficient between the seed time courses and the time courses from each of the voxels in the brain. Additionally, ROI-to-ROI correlation values were obtained by computing correlation coefficients between the pIns seed region and a ROI corresponding to Broadmann Area 31 (BA31), which includes the PCC.

To assess whether pIns-to-PCC connectivity was altered in relation to perceived pain intensity, a Spearman’s rank order correlation coefficient was computed. This non-parametric test was used because it is robust to outliers. Next, a Kruskal-Wallis test one way ANOVA by ranks was conducted to compare average pIns-to-PCC connectivity values across conditions (i.e., *Rest, Pain, and Tactile Sensation*). Lastly, a receiver operating characteristic (ROC) curve was generated to assess the discrimination performance of pIns-to-PCC connectivity across multiple thresholds. Group average connectivity maps for *Rest, Tactile Sensation*, and *Pain* were generated using a second level mixed-effects analysis in Conn Toolbox. Significant clusters are shown with FDR-corrected threshold of p< 0.05. All images are displayed in neurologic convention, with warm colors representing positive correlations, and cool colors representing anti-correlations.

## Results

### Psychophysical results

No subject in either cohort reported experiencing pain during the rest or touch scans; accordingly, these were assigned a pain rating of 0. In Cohort 1, the median (range) subjective pain ratings for the capsaicin scans was 4.5 (0 to 8). One subject from Cohort 1 did not feel pain with the capsaicin stimulus, and reported pain ratings of 0 throughout. This subject’s capsaicin scan was therefore grouped with the *Tactile Sensation* scans. In Cohort 2, the median (range) subjective pain ratings for the first and second capsaicin scans were 3 (1 to 3) and 7 (4 to 7), respectively. Subjective ratings of the salience of the touch and capsaicin stimuli were collected from subjects in Cohort 2. The median (range) of the salience ratings was 6.25 (3 to 8) for the touch stimuli, and 6.5 (4 to 9) for the capsaicin stimuli. A Wilcoxon rank-sum test revealed no significant difference in salience ratings, Z = 0.162, p = 0.871.

### fcMRI results

A total of 40 scans were obtained (two scans each from 6 subjects in Cohort 1, and four scans each from 7 subjects in Cohort 2). These were divided into 3 categories as described in section 4.2, resulting in a total of 13 *Rest* scans, 11 *Tactile Stimulation* scans, and 16 *Pain* scans. Group average functional connectivity maps using the pIns as the seed region for each category are shown in Figure 1. Panel A shows all regions with significant connectivity to the pIns in the *Rest* scans, Panel B shows the same for the Tactile Sensation scans, and Panel C for the *Pain* scans. The *Rest* map is notable for significant pIns anti-correlations to the PCC/precuneus. The *Pain* map is notable for the lack of the pIns-PCC anti-correlations seen in *Rest*, and for positive connectivity between the pIns and the anterior cingulate cortex (ACC). Other areas of connectivity to the pIns (with the magnitude of correlation indicated by T-max values) are provided in Tables 1-3. Table 1 contains significant connectivity clusters for the *Rest* scans, Table 2 for the *Tactile Sensation* scans, and Table 3 for the *Pain* scans.

**Figure 1.**
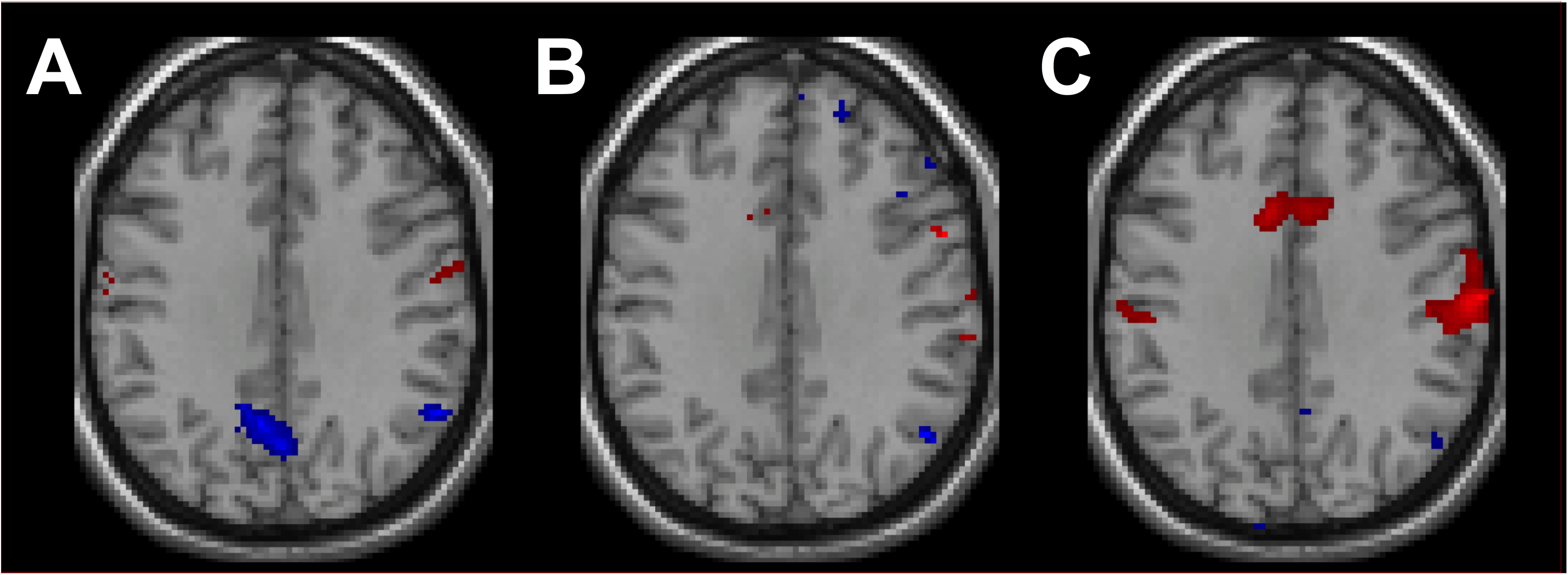
Group average connectivity maps. Maps reflect connectivity to the pIns seed region for *Rest* (panel A), *Tactile Sensation* (panel B), and *Pain* (panel C). Significant correlations are shown in red, and anti-correlations are shown in blue (FDR-corrected p < 0.05). The Rest map demonstrates resting-state anti-correlations between the pIns and the PCC/precuneus. The *Pain* map is notable for the absence of resting state pIns-to-PCC anti-correlations, as well as positive connectivity to the anterior cingulate cortex (ACC).

**Table 1.** Regions with significant connectivity to the pIns in the Rest condition

| Brain Region | Size | T | x, y, z |
| --- | --- | --- | --- |
| Left Insula | 743 | 17.2 | -44, 0, 0 |
| Right Insula | 401 | 13.07 | 40, 0, 10 |
| Precuneus | 230 | -17.04 | -8, -58, 36 |
| Left postcentral gyrus | 135 | 10.37 | -62, 2, 28 |
| Right postcentral gyrus | 105 | 14.34 | 48, -24, 44 |
| Right precentral gyrus | 74 | 10.58 | 62, 10, 8 |
| Right angular gyrus | 66 | -17.16 | 54, -54, 36 |
| Right postcentral gyrus | 59 | 11.05 | 56, -14, 26 |
| Right frontal pole | 54 | -19.44 | 12, 66, 18 |
| Right putamen | 52 | 14.42 | -18, 2, 4 |
| Precuneus | 49 | -11.06 | -2, -44, 44 |
| Right supplementary motor cortex | 46 | 11.15 | -2, 6, 54 |
| Right frontal orbital cortex | 36 | -14.56 | 20, 14, -18 |
| Right frontal pole | 29 | -26.21 | 38, 50, -18 |

Cluster size, max T, and MNI coordinates for regions with significant connectivity to the plns in the Rest condition. Cluster-size threshold $p\text{-FDR} < 0.001$ , height threshold $p < 0.001$ .

**Table 2.**
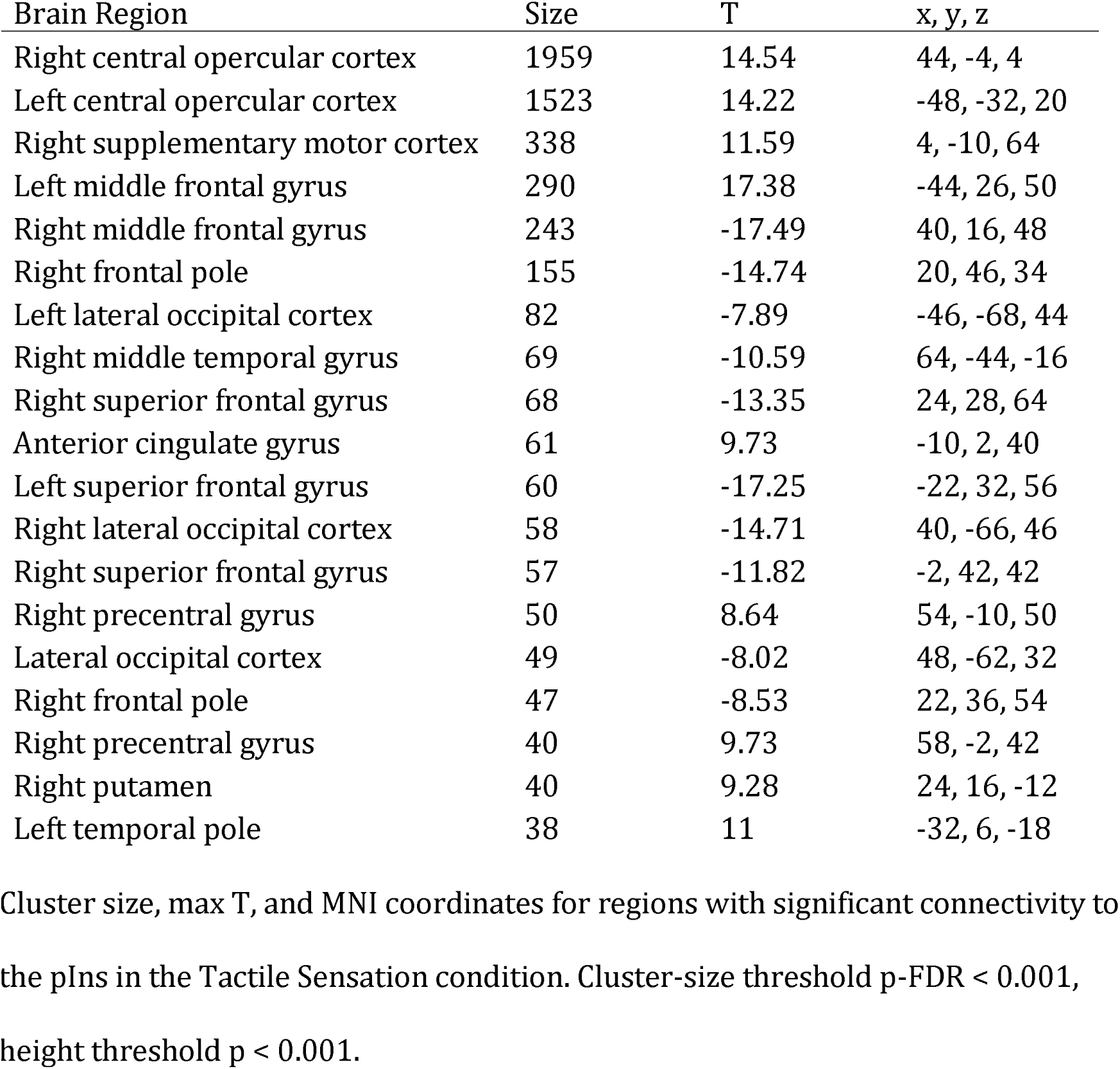
Regions with significant connectivity to the pIns in the Tactile Sensation condition

| Brain Region | Size | T | x, y, z |
| --- | --- | --- | --- |
| Right central opercular cortex | 1959 | 14.54 | 44, -4, 4 |
| Left central opercular cortex | 1523 | 14.22 | -48, -32, 20 |
| Right supplementary motor cortex | 338 | 11.59 | 4, -10, 64 |
| Left middle frontal gyrus | 290 | 17.38 | -44, 26, 50 |
| Right middle frontal gyrus | 243 | -17.49 | 40, 16, 48 |
| Right frontal pole | 155 | -14.74 | 20, 46, 34 |
| Left lateral occipital cortex | 82 | -7.89 | -46, -68, 44 |
| Right middle temporal gyrus | 69 | -10.59 | 64, -44, -16 |
| Right superior frontal gyrus | 68 | -13.35 | 24, 28, 64 |
| Anterior cingulate gyrus | 61 | 9.73 | -10, 2, 40 |
| Left superior frontal gyrus | 60 | -17.25 | -22, 32, 56 |
| Right lateral occipital cortex | 58 | -14.71 | 40, -66, 46 |
| Right superior frontal gyrus | 57 | -11.82 | -2, 42, 42 |
| Right precentral gyrus | 50 | 8.64 | 54, -10, 50 |
| Lateral occipital cortex | 49 | -8.02 | 48, -62, 32 |
| Right frontal pole | 47 | -8.53 | 22, 36, 54 |
| Right precentral gyrus | 40 | 9.73 | 58, -2, 42 |
| Right putamen | 40 | 9.28 | 24, 16, -12 |
| Left temporal pole | 38 | 11 | -32, 6, -18 |

Cluster size, max T, and MNI coordinates for regions with significant connectivity to the plns in the Tactile Sensation condition. Cluster-size threshold $p\text{-FDR} < 0.001$ , height threshold $p < 0.001$ .

**Table 3.** Regions with significant connectivity to the pIns in the Pain condition

| Brain Region | Size | T | x, y, z |
| --- | --- | --- | --- |
| Right Insula | 4598 | 8.21 | 66, -16, 34 |
| Left Insula | 1988 | 7.33 | -44, -6, -2 |
| Anterior cingulate gyrus | 1669 | 8.44 | 2, -10, 62 |
| Postcentral gyrus | 539 | 8.48 | 14, -34, 78 |
| Precuneus | 330 | -7.04 | 6, -66, 48 |
| Lateral occipital cortex | 307 | -7.29 | 30, -74, 48 |
| Right frontal pole | 202 | -6.13 | 36, 60, 0 |
| Left occipital pole | 188 | -6.76 | -4, -98, -6 |
| Right frontal pole | 111 | -9.25 | 14, 66, 14 |

Cluster size, max T, and MNI coordinates for regions with significant connectivity to the plns in the Pain condition. Cluster-size threshold $p\text{-FDR} < 0.001$ , height threshold $p < 0.001$ .

### Association between pIns-to-PCC connectivity and pain

To assess the relationship between pIns-to-PCC connectivity and subjective pain intensity, a Spearman’s rank order correlation coefficient was computed. This was found to be significant, with increased pain ratings being associated with increased pIns-to-PCC connectivity, *r_s_* (38) = 0.34, *p* = 0.032. A scatter plot showing the relationship between connectivity values and subjective pain ratings from all capsaicin scans is shown in Figure 2.

**Figure 2.**
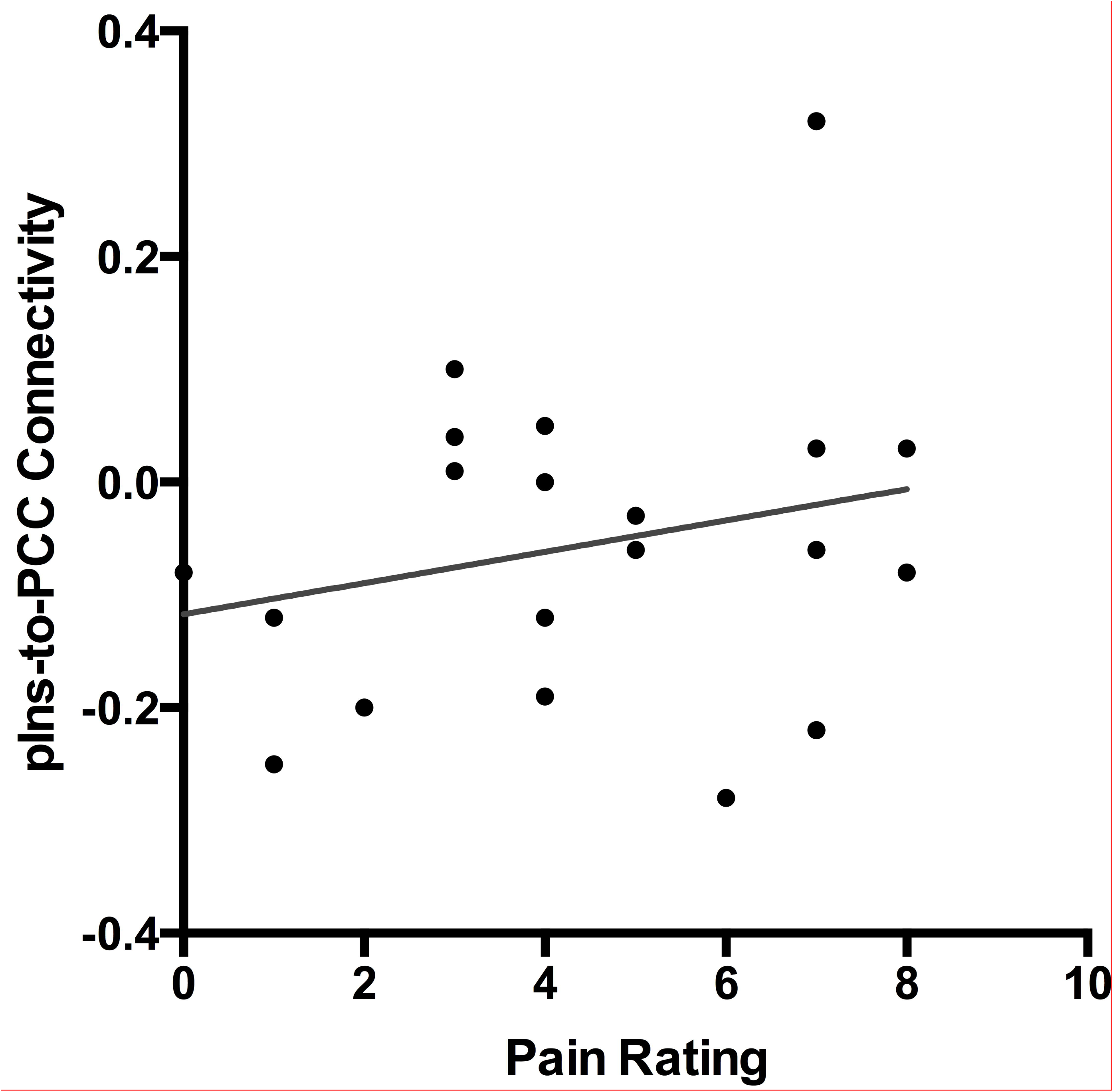
Relationship between connectivity and pain intensity. Scatter plot with linear trendline illustrating the positive correlation between subjective pain ratings and pIns-to-PCC connectivity for all capsaicin scans.

The mean (standard deviation) of pIns-to-PCC connectivity values across conditions were as follows: *Rest* −0.18 (0.15), *Tactile Sensation* −0.10 (0.11), and *Pain* −0.04 (0.14). Note that negative connectivity frequently occurs as a result of global signal regression, and does not necessarily represent “deactivation”. A graphical representation of these values can be found in Figure 3. A Kruskal-Wallis test revealed significant differences in connectivity across conditions, *H*(2) = 9.36, *p* = 0.0093. Post hoc comparisons using t-test, with Bonferroni correction, revealed significant differences in connectivity between the *Rest* and *Pain* scans, *p* = 0.012. Because of the graded pattern of connectivity strength, the differences between the *Rest* and *Tactile Sensation* scans and between the *Tactile Sensation* and *Pain* scans were not statistically significant.

**Figure 3.**
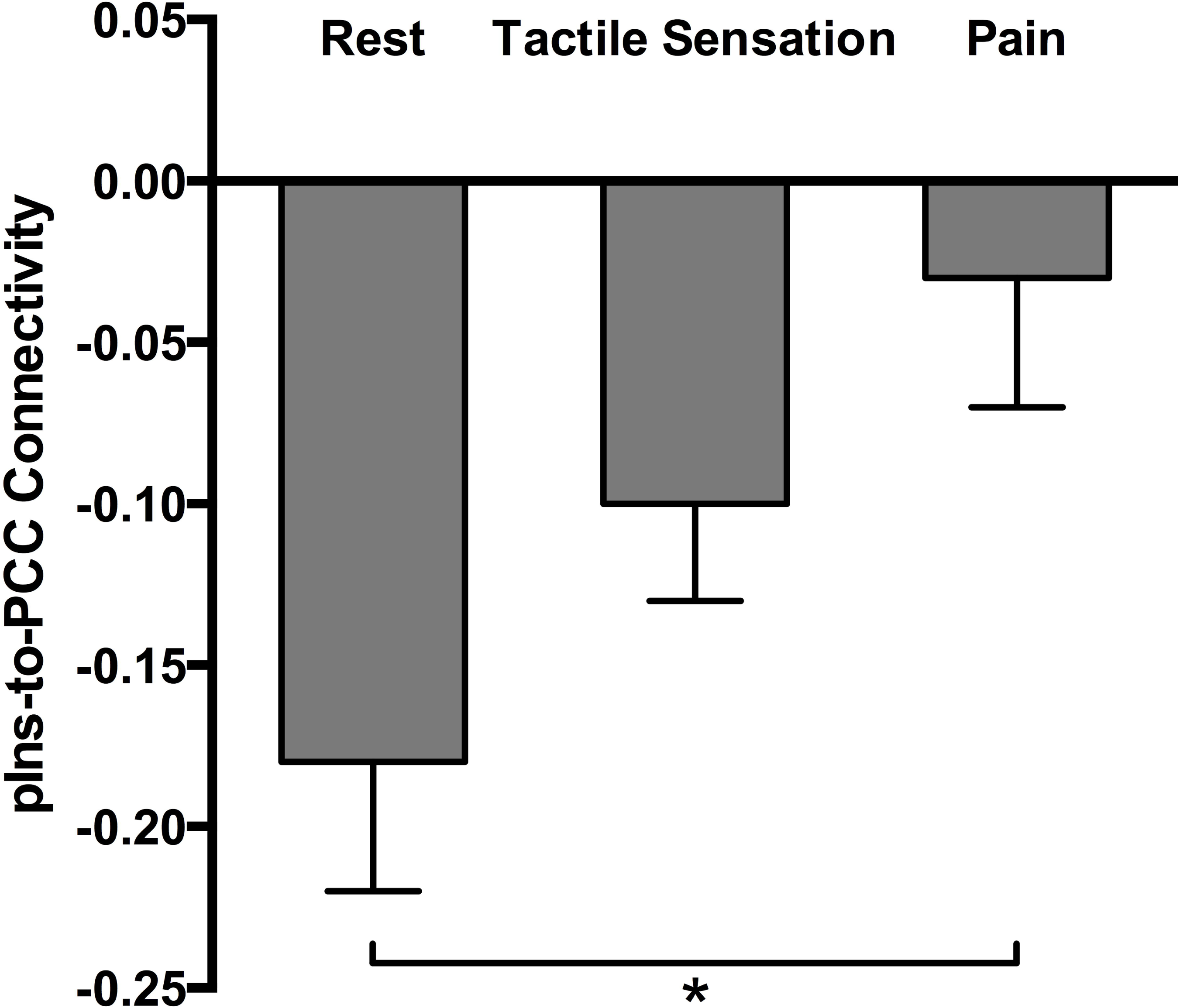
Mean pIns-to-PCC connectivity and standard error by condition. Connectivity was most negative for the *Rest* scans, and least negative for the *Pain* scans. The means were found to significantly differ across conditions, *F*(2,37) = 4.75, *p* = 0.015. Post hoc analyses revealed a significant difference between *Rest* and *Pain*, and not between *Rest* and *Tactile Sensation*; \**p* < 0.05.

Lastly, to assess the overall capacity of pIns-to-PCC connectivity to differentiate *Pain* scans from *Rest* and *Tactile Sensation* scans, a receiver operating characteristic (ROC) curve was generated. The area under the curve was found to be significant at 0.75 (95% CI = 0.59 – 0.91; *p* = .008). This is shown in Figure 4.

**Figure 4.**
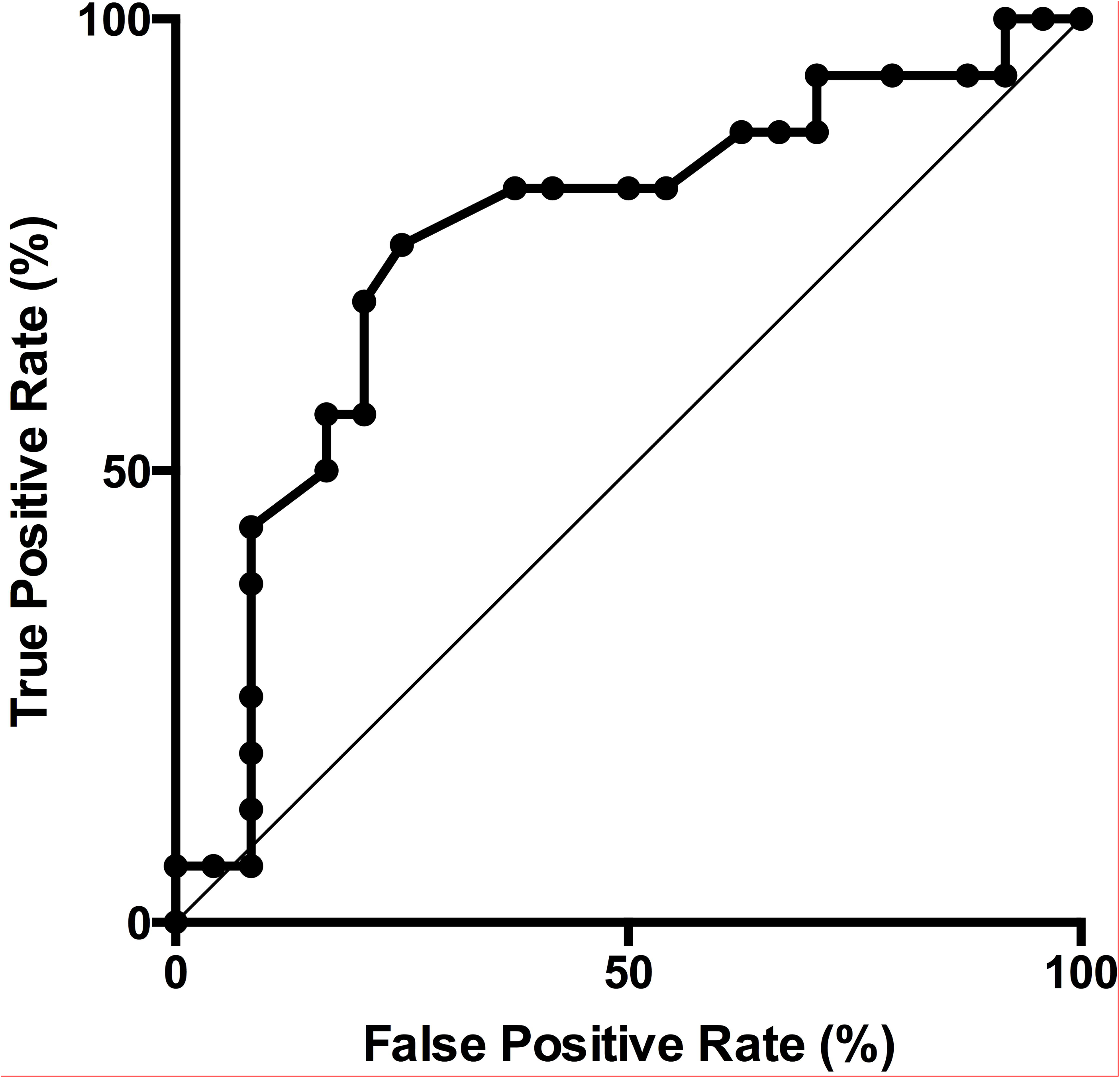
ROC curve for pIns-to-PCC connectivity in classifying pain state. This receiver operating characteristic (ROC) curve shows the classification performance of pIns-to-PCC connectivity in detecting pain across various discrimination thresholds. Area under the curve = 0.75, *p* < 0.01.

## Discussion

Several studies have identified a correlation between insula-to-DMN connectivity and pain intensity in chronic pain patients (Baliki et al. 2014; Loggia et al. 2013; Napadow et al. 2010). We now report a similar relationship between insula-to-DMN connectivity and pain intensity in healthy subjects who are experiencing ongoing acute pain. Additionally, in chronic pain patients, maneuvers that exacerbate chronic pain further alter insula-to-DMN connectivity (Loggia et al. 2013), while pain relief is associated with shifts toward baseline connectivity (Ceko et al. 2015; Napadow et al. 2012). Taken together, these findings point to a relationship between insula-to-DMN connectivity and the magnitude of ongoing pain in both healthy subjects and chronic pain patients. This suggests that the observed connectivity changes in chronic pain patients may be related to ongoing pain rather than more permanent underlying alterations.

Previous studies have identified insula-to-DMN connectivity changes in response to acute pain (Ibinson et al. 2015; Zhang et al. 2014). However, these studies did not attempt to quantify the relationship between pain intensity and changes in connectivity, as was done here. Furthermore, these studies did not include a non-painful control condition. This makes interpretation difficult, as it could be argued that their results simply reflect differences in salience across conditions, rather than pain specific responses. In support of this possible interpretation, there is evidence that patterns of neural activity seen in response to pain largely overlap with patterns seen in response to other salient stimuli (Iannetti and Mouraux 2010; Legrain et al. 2011; Liberati et al. 2016). We included a salient non-painful control condition in an attempt to address this issue, hypothesizing that pIns-to-PCC connectivity would not be altered by salient non-painful stimulation. This was supported by the observation that the rest and tactile stimulation conditions did not significantly differ in terms of pIns-to-PCC connectivity, whereas there was a significant difference between the rest and pain conditions.

Yet it could still be argued that our results simply reflect the greater salience of a painful stimulus relative to a touch stimulus, since connectivity also did not significantly differ between the *Tactile Sensation* and *Pain* conditions and the lack of a difference between *Rest* and *Tactile Sensation* could be attributed to low power. While this cannot be definitively ruled out, the subjective salience ratings we obtained for the touch and capsaicin stimuli did not differ, making an explanation based purely on stimulus salience less likely.

Our secondary aim was to assess the strength of pIns-to-PCC connectivity as a classifier of pain state. This was accomplished through a ROC curve analysis, which showed significant classification accuracy for pIns-to-PCC connectivity in discriminating pain states from non-pain states (i.e., *Rest* and *Tactile Sensation*). Several groups have previously demonstrated the utility of fMRI in detecting pain, though with important limitations (Brown et al. 2011; Marquand et al. 2010). Perhaps the best example comes from Wager et al. (Wager et al. 2013), who used task-based fMRI and machine learning to develop a model capable of accurately differentiating pain from warmth and other aversive events. While this represents a significant achievement, the potential applications of this approach are limited by the reliance on a block-design paradigm, which requires the repeated presentation of an experimental stimulus. In contrast, a functional connectivity-based approach would not necessarily require an experimental stimulus. Such a technique would be better suited to measure ongoing pain, as would be seen in clinical populations. The present findings suggest that such an fcMRI-based approach may be possible.

The posterior insula was the focus of the present study. There are several lines of evidence to suggest that this region plays a key role in pain processing. First, anatomical studies in primate models have revealed projections from pain sensitive thalamic nuclei to the posterior insula (Burton and Jones 1976; Friedman and Murray 1986; Mufson and Mesulam 1984). In fact, the posterior insula has been shown to receive more spinothalamic input than any other cortical area (Dum et al. 2009). Second, consistent with these anatomical connections, the insula is the region most commonly activated in response to pain across imaging studies (Apkarian et al. 2005). Furthermore, Segerdahl et al. (Segerdahl et al. 2015) found that blood flow in the pIns was correlated with induced fluctuations in pain intensity over time. Third, lesion studies show that damage to the posterior insula leads to increased pain thresholds (Birklein et al. 2005; Bowsher et al. 2004; Greenspan and Winfield 1992; Kim 2007; Schmahmann and Leifer 1992), while lesions to the anterior insula do not (Greenspan et al. 1999). Posterior insular lesions are also implicated pain asymbolia, a condition where patients demonstrate no primary sensory defects, yet fail to show typical emotional and withdrawal responses to painful stimuli (Berthier et al. 1988). Finally, electrical stimulation of the posterior insula and adjacent parietal operculum can evoke painful sensations (Isnard et al. 2004; Mazzola et al. 2009; Ostrowsky et al. 2002). While only a minority of stimuli applied to this region cause pain, no other cortical region has been shown to give rise to pain when electrically stimulated (Mazzola et al. 2012). A case has also been reported of a patient with purely painful seizures who was found to have an epileptogenic focus in the posterior insula, further suggesting that the experience of pain can be generated by neural activity originating in this region (Isnard et al. 2011). In sum, the posterior insula shares anatomical connections with ascending pain pathways, shows blood flow that is correlated with perceived pain intensity, leads to decreased pain sensitivity when damaged, and gives rise to pain when stimulated. While it should be noted that this region is involved in a wide range of functions (Nieuwenhuys 2012), these findings nonetheless suggest an important role for the posterior insula that is specific to pain processing. Thus, it is unsurprising that a measure based on connectivity of this region would be useful in the detection of pain. In fact, the pIns was among the regions assigned the highest predictive weight in the Wager et al. (2013) pain detection model.

Similarly, the DMN seems to be characteristically affected by pain. Baliki et al. (Baliki et al. 2014) examined the effects of several chronic pain conditions on multiple resting state networks (RSNs), and found the DMN to be the only RSN consistently altered across chronic pain conditions. The authors also noted corresponding changes in insula-to-DMN connectivity in association with chronic pain. Similar alterations have been reported in a number of chronic pain conditions including fibromyalgia (Napadow et al. 2010), chronic back pain (Tagliazucchi et al. 2010), complex regional pain syndrome (Baliki et al. 2014), diabetic neuropathic pain (Cauda et al. 2009), and osteoarthritis (Baliki et al. 2014). Moreover, Ichesco et al. (Ichesco et al. 2014) specifically found altered pIns-to-PCC connectivity in fibromyalgia patients compared to controls.

As noted above, the current study is limited by the use of a relatively small sample and the reliance on a single pain stimulus. It is worth noting that our findings are consistent with results obtained using an electrical pain stimulus in healthy subjects (Ibinson et al. 2015), and, as noted above, are broadly consistent with studies involving chronic pain patients. Additionally, we collected data using different parameters on two separate scanners, but still found that the relationship between connectivity and pain intensity persisted. This tempers concerns about the methodology and adds to the robustness of these findings.

## Conclusions

In summary, we have demonstrated that ongoing pain in healthy subjects causes alterations in posterior insula-to-DMN connectivity in proportion to perceived pain intensity. These findings are not explained entirely by salience. Our results share similarities with previous reports of connectivity changes in chronic pain patients, yet we expand on this prior work by demonstrating corresponding alterations in response to acute pain in healthy subjects. Furthermore, pain state can be classified based on pIns-to-PCC connectivity.. These findings suggest that, with further development, fcMRI measures based in part on pIns-to-PCC connectivity may hold promise as a noninvasive biomarker for pain.

## Declarations

### Ethics approval and consent to participate

Institutional Review Board approval was obtained from the University of Pittsburgh, and informed consent was obtained from each subject. All procedures performed in studies involving human participants were in accordance with the ethical standards of the institutional and/or national research committee and with the 1964 Helsinki declaration and its later amendments or comparable ethical standards.

### Consent for publication

not applicable

### Availability of data and material

The datasets used and/or analyzed during the current study are available from the corresponding author on reasonable request.

### Authors’ contributions

CB, KV, and JI all made substantial contributions to conception, design, acquisition of data, analysis and interpretation of data, and drafting and revising the manuscript.

## Acknowledgements

Not applicable

